# Characterization and Antiatherogenic Potential of P3R99 Monoclonal Antibody Against Sulfated Glycosaminoglycans: Physicochemical and Functional Insights

**DOI:** 10.1101/2024.09.09.612153

**Authors:** Leidy M. Valencia, Yoandra Martínez-Montano, José A. Gómez, Roger Sarduy, Arletty Hernández, Spencer Proctor, Aymé Fernández-Calienes, Víctor Brito, Yosdel Soto

**Affiliations:** Department of Immunobiology, Centre for Molecular Immunology, Havana, Cuba; Department of Quality Control, Centre for Molecular Immunology, Havana, Cuba; Metabolic and Cardiovascular Diseases Laboratory, Division of Human Nutrition, Alberta Diabetes Institute, University of Alberta, Edmonton, Alberta, Canada; Innovation and Managing Direction, Centre for Molecular Immunology, Havana, Cuba

**Author notes:** **Correspondence:** Dr. Yosdel Soto, Department of Immunobiology, Centre for Molecular Immunology, Havana, Cuba.

**Keywords:** atherosclerosis, immunotherapy, monoclonal antibodies, proteoglycan, protein expression, recombinant protein characterization

## Abstract

Atherosclerosis is initiated by the retention of ApoB-containing lipoproteins in the arterial wall, mediated by glycosaminoglycan chains of proteoglycans. At the Center for Molecular Immunology, we are developing the P3R99 monoclonal antibody (mAb) to target this process. This study characterizes new P3R99 mAb variants expressed in CHO-K1 and HEK-293 cell lines. We compared these variants with the parental mAb from NS0 cells using SDS-PAGE, size exclusion and cation exchange chromatography, dynamic light scattering, peptide mapping, far-UV circular dichroism, and PNGase F deglycosylation. All variants exhibited a molecular size of ∼150 kDa, ∼99% purity, and similar average particle sizes (12.5-13.7 nm). They displayed a high β-sheet content (>40%) and basic amino acids on the surface, with minor differences in peptide maps compared to the parental mAb. Notable differences were found in the content of acidic and basic species and glycosylation profiles. NS0-derived P3R99 had lower G0F content (10.39%), higher G1F (38.29%) and G2F (30.44%) levels, with more terminal galactose (83.07%) and sialylation (15.33%). In contrast, CHO-K1 and HEK-293 variants showed similar glycosylation patterns. Despite these differences, the antigen and atherosclerotic lesion recognition properties of the mAb were unaffected *in vitro*. Biodistribution studies in Sprague Dawley rats (1 mg, IV, n=3) revealed preferential accumulation of the new P3R99 variants in aortas and reduced LDL arterial retention (1 mg, IP). Passive administration of the mAbs (2 mg every three days, three IV doses, n=6-7) in a Lipofundin 20%-induced atherosclerosis NZW rabbit model also demonstrated preferential accumulation in aortas and reduced atherosclerosis, with 60% of treated rabbits not developing lesions. These results suggest that the P3R99 mAb derived from CHO-K1 and HEK-293 cells retains its antiatherogenic properties despite structural differences from the NS0-derived mAb associated with the different expression systems.

## INTRODUCTION

Cardiovascular diseases related to atherosclerosis remain the leading cause of mortality worldwide (Gaidai et al., 2023). Atherosclerosis is a chronic inflammatory disorder of the arterial wall characterized by the focal deposition of lipids (atherosis) and fibrous tissue (sclerosis) in the intimal layer of medium and large-caliber arteries, including the aorta and coronary arteries (Andersson et al., 2010). A critical early event in this process is the retention of ApoB-containing lipoproteins such as low-density lipoprotein (LDL), lipoprotein (a), and triglyceride-rich lipoproteins in the subendothelial space (Williams and Tabas 1995; Tabas et al., 2007; Tannock and King 2008; Fogelstrand and Boren 2012). This retention is mediated by interactions between positively charged residues of ApoB and negatively charged sulfate groups on glycosaminoglycan (GAG) side chains of arterial proteoglycans space (Williams and Tabas 1995; Borén et al., 1998; Skålén et al., 2002; Tabas et al., 2007). The retention process leads to lipoprotein modifications that activate endothelial cells and macrophages, ultimately contributing to foam cell formation and inflammatory signaling (Hansson and Hermansson 2011).

Current therapeutic approaches for atherosclerosis primarily target risk factors such as hypercholesterolemia, hypertension, and diabetes (Amir and Binder 2010) and aim to modulate lipid metabolism and subendothelial inflammation (Björkbacka et al., 2013; Bäck and Hansson 2015; Ladeiras-Lopes et al., 2015). However, interventions targeting vascular wall components have been less explored, with only a few candidates documented in preclinical phases (Zeng et al., 2005; Ballinger et al., 2010; Kumarapperuma et al., 2024). The Center for Molecular Immunology (CIM, Havana, Cuba) has developed an innovative immunotherapy targeting subendothelial lipoprotein retention using the chimeric monoclonal antibody (mAb) P3R99 (Soto et al., 2012; Soto et al., 2024). P3R99 specifically recognizes GAGs, particularly chondroitin sulfate (CS), and inhibits the association of LDL and chylomicron remnants with CS proteoglycans (CSPGs), thereby preventing lipoprotein retention and subsequent oxidative modifications in arterial walls (Soto et al., 2012; Soto et al., 2024)

The P3R99 mAb (hereafter referred to as R99) is highly immunogenic, with idiotypic regions that induce antibodies with similar specificity *in vivo*, thereby conferring atheroprotective effects through active immunization (Brito et al., 2012; Soto et al., 2012; Sarduy et al., 2017; Soto et al., 2024). Immunization with R99 in animal models of atherosclerosis prevents lesion formation and halts or reduces the progression of pre-existing lesions, particularly at higher doses where it promotes regression of advanced lesions (Brito et al., 2017; Sarduy et al., 2017). These effects are linked to the induction of anti-anti-idiotype antibodies (Ab3) that mimic the specificity of R99, inhibiting the subendothelial retention of proatherogenic lipoproteins (Brito et al., 2012; Soto et al., 2012; Soto et al., 2024). Despite the well-documented active immunization effects, the potential of R99 for direct blocking (passive effect) in disease prevention and progression has yet to be fully explored.

R99 was originally expressed in NS0 mouse myeloma cells; however, the choice of host cell lines, transfection systems, and culture conditions are critical for the efficient production of recombinant proteins in the biopharmaceutical industry (Chusainow et al., 2009). Mammalian cells, such as Chinese hamster ovary (CHO-K1) and human embryonic kidney (HEK-293) cells, are widely used in the production of complex molecules like mAbs due to their scalability, high-density growth in suspension cultures, use of serum-free and animal component-free media, appropriate protein folding, and human-like glycosylation pattern (Fernández and Vega 2016). R99 mAb is also expressed in CHO-K1 and HEK-293 cell lines preadapted to serum-free media resulting in two new variants of this mAb: CHO-R99 and HEK-R99.

In this study, we aimed at conducting a comprehensive physicochemical characterization of the R99 mAb expressed in NS0, CHO-K1, and HEK-293 cells to evaluate how the expression system affects the structural integrity, stability, and functional properties of R99 mAb. A key focus of this study was validating their recognition properties, its biodistribution *in vivo*, along with evaluating their antiatherogenic properties as a passive immunotherapy, providing insights into the mechanistic actions and therapeutic implications of targeting GAGs in cardiovascular disease.

## MATERIALS AND METHODS

### Antibodies

NS0-R99 (Soto et al., 2012), CHO-R99, HEK-R99 and humanized R3 (hR3, anti-human epidermal growth factor receptor) (Mateo et al., 1997) were purified from the cell culture supernatants by affinity chromatography using Protein A-MabSelect SuRe (GE Healthcare, USA). Purity was assessed by SDS-PAGE under both non-reducing and reducing conditions, following standard protocols (Laemmli 1970). The EO6 mAb, specific for oxidized phospholipids, was purchased from Avanti Polar Lipids (Alabaster, AL).

### Lipofundin MCT/LCT 20%

Lipofundin medium-chain triglycerides/short chain triglycerides 20% (hereafter referred to as Lipofundin 20%) was acquired from Braun (Melsungen AG, Germany). Lipofundin is a lipid emulsion containing 100 g of soybean oil, 100 g of medium-chain triglycerides, 25 g of glycerol, 12 g of egg lecithin, 170 ± 40 mg of α-tocopherol, and sodium oleate in water for injection, sufficient to make up 1000 mL.

### Animals

Sprague-Dawley male rats (241-260 g, 6 weeks old) and New Zealand white (NZW) female rabbits (1.8-2.0 kg, 12 weeks old) were obtained from the *Centro Nacional para la Producción de Animales de Laboratorio* (CENPALAB, Mayabeque, Cuba). Animals were housed at the Centre for Molecular Immunology (Havana, Cuba) under standard conditions (25°C, 60±10% humidity), 12 h day/night cycles with water and food *ad libitum.* All procedures were approved by the Institutional Animal Ethic Committee, following its animal care and use guidelines.

### Size Exclusion Chromatography

Size exclusion chromatography (SEC-HPLC) was performed using an LC-2030C chromatograph (Shimadzu, Japan) equipped with a Yarra™ SEC 3000 column (Phenomenex, CA), according to manufacturer’s instructions. The mobile phase consisted of 8.1 mM Na_2_HPO_4_, 1.5 mM KH_2_PO_4_, and 0.4 M NaCl in 1 L of purified water, adjusted to pH 7.2–7.4. Samples were analyzed at a flow rate of 1 mL/min, with the column maintained at 25°C and the system operated at a maximum pressure of 10 MPa. Chromatographic separation was achieved isocratically, and protein elution was monitored by UV detection at 280 nm.

### Dynamic Light Scattering

Particle size distribution of the samples was determined using a dynamic light scattering (DLS) Delsanano C particle analyzer (Beckman Coulter, IN), employing a 658 nm diode laser at 30 mW. A 100 µL sample was placed in a glass cuvette and analyzed at 25°C with a scattering angle of 165°. Data acquisition involved triplicate measurements of 100 counts each, utilizing Cumulant and Contin statistical methods (Gun’ko et al., 2003).

### Peptide Mapping

Samples underwent digestion following denaturation and reduction with 6 M guanidine hydrochloride, 2.38 M Tris, and 500 mM dithiothreitol at 37°C for 1 h. Alkylation was performed using 375 mM iodoacetamide for 20 min in the dark. Thereafter, samples were desalinated with 0.1 M (NH_4_)HCO_3_ (pH 7.5) using pre-equilibrated Sephadex™ G-25 DNA Grade columns (Cytiva, Uppsala, Sweden). Digestion was conducted overnight at 37°C with a trypsin/Lys-C enzyme mixture (1:50 ratio). The reaction was stopped with trifluoroacetic acid, and peptides were analyzed by RP-HPLC (Shimatzu, Kyoto, Japan) on an AdvanceBio Peptide Plus C18 column (Agilent Technologies, Santa Clara, CA). The samples (100 µL) were injected at a flow rate of 0.8 mL/min, under a pressure of 30 MPa, and at 55°C. Mobile phases consisted in purified water with 0.1% trifluoroacetic acid and acetonitrile with 0.1% trifluoroacetic, respectively. Peptides were detected at 214 nm using an LC-2030C chromatograph integrated with LabSolutions Version 5.93 for method optimization and data processing (Jiang et al., 2020).

### Cation Exchange Chromatography

Cation exchange chromatography was conducted using an LC-2030C chromatograph with a MAbPac SCX column (Thermo Scientific, Whaltham, MA). Mobile phase A comprised 20 mM 2-(N-morpholino) ethanesulfonic acid, 1 M NaCl in 500 mL purified water (pH 5.6), while mobile phase B contained 20 mM 2-(N-morpholino) ethanesulfonic acid, and 60 mM NaCl in 2 L purified water (pH 5.6). Samples were analyzed at a flow rate of 0.7 mL/min, at 30°C, and a maximum pressure of 32 MPa. Samples (50 µg) were injected, and detection was performed at 280 nm. Each of the three R99 variants (NS0, CHO, HEK) was run in duplicate. Additionally, samples treated with carboxypeptidase B were analyzed to exclude the presence of terminal lysine residues. Chromatograms were manually integrated, and the area percentages of acidic, basic, and main peaks were calculated.

### Enzymatic Deglycosylation with PNGase F

Deglycosylation was performed on 100 µg of each R99 lot, adjusted to 90 µL with purified water, using a PNGase F kit devoid of glycerol at 37°C for 2 h. Then, proteins were precipitated by adjusting to room temperature, mixing with three volumes of cold absolute ethanol, and incubating at −20°C for 2 h. After centrifugation at 10,000 rpm for 10 min at 4°C, the supernatants were collected and dried in a vacuum evaporator. Resuspended samples in 5 µL of 2-AB labeling solution, consisting in 2 M 2-aminobenzamide and 1 M NaCNBH_3_ in DMSO: acetic acid (7:3, v/v), were incubated at 65°C for 2 h. For sample cleanup, Sephadex™ G-25 DNA Grade columns equilibrated with purified water, 30% acetic acid, and acetonitrile were used. Glycans were eluted with purified water, dried in a vacuum, and stored at −20°C. Chromatographic analysis was performed on a modular chromatograph with a TSKgel Amida-80 column (Tosoh Bioscience, Tokyo, Japan), using an 80-47% acetonitrile gradient in 50 mM ammonium formate over 167 minutes, with a flow rate of 0.4 mL/min and detection by fluorescence (λ_exc = 330 nm, λ_det = 420 nm).

### Far-UV Circular Dichroism Spectroscopy

Far-UV CD spectra were acquired with a Jasco V-730 Spectrophotometer (Jasco Corporation, Tokyo, Japan), calibrated with d-10-camphorsulfonic acid and equipped with a temperature controller and mini-circulating water bath. Measurements were performed in 0.1 cm quartz cuvettes over a range of 190-250 nm, with a 0.1 nm interval, a scanning speed of 20 nm/min, and a response time of 8 seconds. Spectra were analyzed using Spectra Manager-v2 software (Jasco) with baseline correction, blank subtraction, and smoothing (Adaptive-Smoothing). Molar ellipticity per residue ([θ]λ) was calculated as: [θ]λ=100⋅θλ/(C⋅n⋅L), where θλ represents the ellipticity in mdeg, n is the number of amino acid residues, C is the protein concentration in mmol/L, and L is the path length in cm. Secondary structure content was estimated by deconvolution of FAR-UV CD spectra using the BeStSel server (Micsonai et al., 2018; Micsonai et al., 2021)(Woody 1996).

### Solid phase assays

#### Quantification of human IgG

Human IgG levels in serum and tissue homogenates were quantified by an Enzyme-Linked Immunosorbent Assay (ELISA), with appropriate sample dilutions standardized for each specimen. High-binding 96-well microtiter plates (Costar, Cambridge, MA) were coated overnight at 4°C with a goat anti-human IgG Fc antibody (Sigma) at 3 µg/mL in Carbonate-Bicarbonate buffer (0.1 M, pH 9.6). After washing with Phosphate-Buffered Saline (PBS, 137 mM NaCl, 2.7 mM KCl, 10 mM Na_2_HPO_4_, 1.8 mM KH_2_PO_4_) containing 0.05% Tween-20, plates were blocked with 1% bovine serum albumin (BSA)/PBS for 1 h at room temperature. The samples and standard human IgG (7.81-500 ng/mL) diluted in blocking solution were incubated for 2 h, followed by the addition of a goat anti-human IgG Fc-alkaline phosphatase conjugate (Sigma). The reaction was developed using paranitrophenylphosphate substrate (1 mg/mL) in Diethanolamine buffer (1M diethanolamine, 0.5 M MgCl_2_-6H_2_O, pH 9.8) in an ELISA reader (Dialab ELx800, Vienna, Austria).

#### Recognition of R99’s anti-idiotype antibody 1E10

The ability of the new R99 mAb variants to recognize the anti-idiotypic mAb 1E10 was evaluated by ELISA under conditions similar to those described above. Maxisorp plates (Nunc) were coated with 1E10 mAb at a concentration of 10 µg/mL. The plates were then incubated with 50 µL/well of mAbs derived from different cell lines (1.5-100 ng/mL). Detection was carried out using a goat anti-human IgG Fc-specific antiserum conjugated to alkaline phosphatase (Sigma-Aldrich, St. Louise, MO), with specificity assessed using the isotype control mAb hR3.

#### Recognition to chondroitin sulfate

The binding of the new R99 mAb variants to CS was assessed following the method previously described by our group (Soto et al., 2012). Maxisorp microtiter plates were coated with CS (Sigma) at 10 µg/mL in HEPES-buffered saline (SSTH, 20 mM HEPES, 150 mM NaCl, pH 7.4) and incubated overnight at 4°C. After three washes with SSTH, the plates were blocked with SSTH containing 1% BSA for 1 h at room temperature. The R99 mAb, derived from different cell lines, was added at concentrations ranging from 0.15-10 µg/mL, with hR3 mAb used under similar conditions as an isotype control. Samples were prepared in binding buffer (10 mM HEPES, 20 mM NaCl, 2 mM CaCl_2_, 2 mM MgCl_2_, pH 7.4) and incubated for 1 h at room temperature. After three washes, a goat anti-human IgG Fc-specific antiserum conjugated to alkaline phosphatase (Sigma) was added, and the absorbance was measured at 405 nm.

#### Induction of anti-idiotypic and anti-CS responses

The anti-idiotypic (Ab2) and anti-anti-idiotypic (Ab3, specific for CS) responses in NZW rabbits were evaluated by ELISA, as previously described by our group (Brito et al., 2012; Soto et al., 2024). To assess the Ab2 response, sera collected on the day of sacrifice were tested for reactivity against R99 and hR3 mAbs. Reactivity to CS was similarly evaluated in both pre-immune and post-immune sera. In both cases, a goat polyclonal antiserum specific for rabbit IgG conjugated to horseradish peroxidase (Jackson ImmunoResearch Lab, West Grove, PA) was used, and absorbance was measured at 490 nm. Sera from seven rabbits previously immunized with NS0-R99 mAb (100 µg weekly, 4 weeks, SC) were used as immunogenicity controls.

### Studies in rats

The accumulation of R99 mAb variants in the aorta and their effects on LDL retention/oxidation were assessed in rats as previously described (Calara et al., 1998; Soto et al., 2012). Rats received a 1 mg bolus IV injection of either R99 mAb variants or the hR3 control, followed by LDL administration 1 h later (1 mg, IP) (Figure 3A). The mAb dose was approximately 4 mg/kg, equivalent to 270 mg in humans, based on standard allometric dose conversion methods (Nair et al., 2016; Germovsek et al., 2021). A control group receiving only 1 mL PBS IV was also included. After 24 h, rats were euthanized, the aortas were excised, and the aortic arch was fixed in 4% formaldehyde at 4°C for 24 h before embedding in paraffin. Blood was drawn at the end of the experiment and additional tissues, including the heart, liver, lung, kidney, and aorta segments, were collected and stored at −80°C for further analysis.

### Experiments in NZW rabbits

Atherosclerosis was induced in NZW female rabbits by administering Lipofundin 20% through the marginal ear vein at a dose of 2 mL/kg daily for eight consecutive days (Jellinek et al., 1982; Delgado Roche et al., 2012; Medina et al., 2012). To evaluate the antiatherogenic effects of the new R99 mAb variants *in vivo*, three IV injections of 2 mg of the mAbs were administered through the marginal ear vein every 3 days, concurrently with the induction of the disease (Figure 4A). The study included treatments with CHO-R99 (n=6) and HEK-R99 (n=7) mAbs, as well as a control group receiving an equivalent dose of the mAb R3h (n=6) to assess specificity. In total, 6 mg of mAbs were administered throughout the study (approximately 3 mg/kg), corresponding to an estimated human equivalent dose of 68 mg (Nair et al., 2016; Germovsek et al., 2021). The rabbits were anesthetized with ketamine hydrochloride (5 mg/kg, IM) and euthanized 24 h after the final Lipofundin 20% administration with an overdose of sodium pentobarbital (90 mg/kg, IV)(Abbott Laboratories, Mexico SA de CV, Mexico). Tissue collection included the heart, muscle, liver, lung, kidney, and thoracic and abdominal segments of the aorta, which were stored at −80°C until further analysis. Blood samples were drawn from the marginal ear vein at the start of the study and by intracardiac puncture at the time of sacrifice. In a separate set of experiments, two rabbits were administered Lipofundin 20% to evaluate the recognition of atherosclerotic lesions by the new R99 mAb variants by immunohistochemical studies. In both cases, the aortic arch was excised and embedded in paraffin for subsequent histopathological and immunohistochemical analysis.

### Biodistribution studies of mAb R99

The biodistribution of R99 in rabbits and rats was evaluated by quantitative ELISA in homogenates from tissues collected. Briefly, the organs were macerated using a tissue homogenizer (Miltenyi Biotec, Bergisch Gladbash, Germany) with PBS containing 1.5% Triton X-100 and 1% protease inhibitor cocktail (Sigma). The lysates were centrifuged at 10,000 rpm for 20 min at 4°C. Total protein content was measured using the bicinchoninic acid method (Smith et al., 1985), and the supernatants were stored at −80°C until further analysis. The human IgG content in these organs, as well as the circulating levels of the mAbs at sacrifice, were quantified using the ELISA method described above, with each sample diluted appropriately.

### Histopathological findings

Arterial tissue sections from rabbits were deparaffinized in xylene and rehydrated through decreasing concentrations of ethanol. Histopathological analysis included conventional hematoxylin-eosin (H&E) staining to visualize atheromatous lesions, and Alcian Blue staining (Ström et al., 2004) to detect GAGs in the arterial wall. Verhoeff-Van Gieson staining was used to determine the intima thickness (Ruengsakulrach et al., 1999; Ström et al., 2004; Medina et al., 2012). Sections were analyzed using an Olympus BX51 fluorescence microscope with a DP20 digital camera (Tokyo, Japan). For morphometric analysis, five sections from each rabbit were analyzed using ImageJ 1.52a (National Institute of Health, Bethesda, MD, USA) to measure the intimal thickness and choose the five largest thickness measurements per animal. Also, lesions were classified as mild, moderate, and advanced, based on their size, extracellular matrix content and the presence of extracellular lipids (Suresh et al., 2020).

### Immunohistochemical analysis

Immunohistochemical evaluations were performed on paraffin-embedded aortic arch tissues from rats and rabbits, as described previously (Soto et al., 2012; Soto et al., 2014). The sections were first deparaffinized, rehydrated, and treated to block endogenous peroxidase activity (Dako, Ontario, Canada). For rats, oxidized LDL was detected using EO6 mAb at 20 µg/mL for 30 min at room temperature, followed by biotin-conjugated goat anti-mouse immunoglobulin polyclonal antiserum (Jackson). To assess the reactivity of R99 mAb variants to atherosclerotic lesions, rabbit tissues were incubated with R99 mAbs or the hR3 control at 20 µg/mL for 1 h, followed by biotin-conjugated goat antiserum specific for human IgG (Jackson). The reaction was developed using a ready-to-use diaminobenzidine substrate solution (Dako, Ontario, Canada) for 5 min. Analysis was performed using an Olympus BX51 fluorescence microscope.

### Statistical analysis

The data is expressed as means with standard errors and analysed using GraphPad Prism 11 (GraphPad Software, Inc). All the experiments *in vitro* were conducted at least twice. Reactivity in ELISAs was defined when OD≥0.25 and at least twice the value of the blank or the isotype control reactivity. To analyze the data, the normality of the distribution of the variables was first checked using the Shapiro-Wilk test and the homogeneity of variances with the Bartlett-Box test. For normally distributed data with equal variances, a one-way ANOVA followed by Tukey’s test was used. For not parametric analysis multiple groups comparisons were performed by Kruskal-Wallis’ test followed by Dunn’s multiple comparison. Values of *P* < 0.05 were considered statistically significant and the significance levels are shown in the figures and legends.

## RESULTS

### R99 mAb expressed in different host cell lines exhibited similar structural integrity

We conducted a comprehensive physicochemical characterization of the R99 mAb derived from NS0, CHO-K1, and HEK-293 cell lines to assess their primary and secondary structure, charge profiles, and glycosylation patterns. SDS-PAGE under non-reducing conditions showed a band around 150 kDa for all variants, consistent with the expected molecular weight of a human IgG (Figure 1, left panel). Under reducing conditions, bands at approximately 25 and 50 kDa confirmed the presence of both light and heavy chains, indicating successful expression of R99 with minimal degradation (Figure 1, right panel). SEC-HPLC analysis revealed high purity (>99%) for all variants and low aggregation levels, confirming the overall quality of the mAbs (Not shown). DLS measurements showed that NS0-, CHO- and HEK-derived R99 mAbs were monodisperse, with particle diameters of 13.7, 12.5 and 13.7 nm, respectively (Supplemental Figure 1).

**Figure 1.**
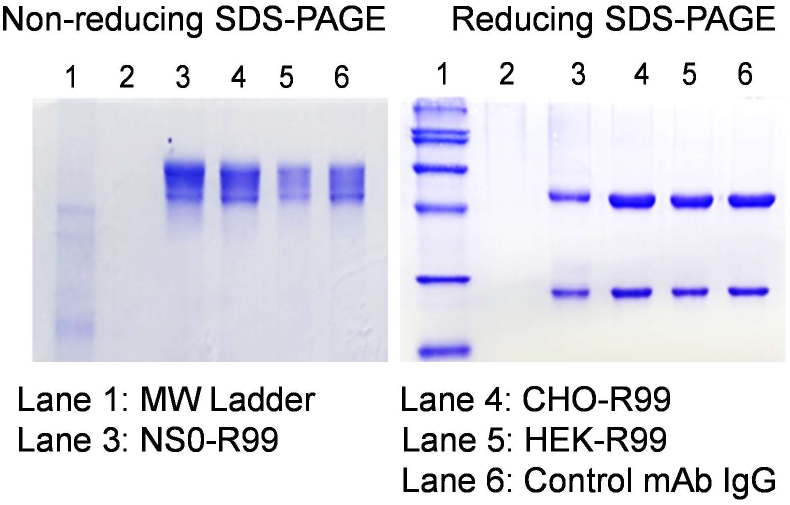
SDS-PAGE analysis of R99 mAb expression in NS0, CHO-K1, and HEK-293 cell lines. SDS-PAGE profiles of R99 mAb produced in NS0, CHO-K1, and HEK-293 cells under non-reducing and reducing conditions. Under non-reducing conditions, all variants displayed a prominent band around 150 kDa, consistent with the expected molecular weight of a complete human IgG, along with minor bands indicating the presence of different isoforms. Under reducing conditions, the expected bands for the heavy chain (∼50 kDa) and light chain (∼25 kDa) were observed, confirming successful expression and proper assembly of the mAb across all host cell lines.

### R99 mAb from different host cells exhibited comparable primary and secondary structure

RP-HPLC-based peptide mapping was performed to assess amino acid sequence similarity and potential post-translational modifications (PTMs) across variants (Judd 2002). Despite similar chromatograms, the parental NS0-R99 mAb displayed 99 tryptic species; while CHO-R99 and HEK-R99 each showed 90 peaks (Supplemental Figure 2). Of these, 89 peaks were common across all variants, 10 were unique to NS0-R99, and one was found in both CHO- and HEK-derived R99 but was barely detectable in the parental mAb. Those differences likely represent variations in glycosylation and other PTMs (Sandra et al., 2018). CHO- and HEK-derived variants showed similar peak compositions, consistent with findings indicating minimal differences when using similar cell lines (Goh and Ng 2018; Duivelshof et al., 2019). Subsequently, far-UV circular dichroism (CD) spectroscopy was used to analyze secondary structure, revealing that all R99 variants exhibit nearly identical spectra with a maximum absorption at around 201 nm and a minimum at 216 nm (Supplemental Figure 3). The secondary structure analysis confirmed that R99 predominantly maintains a β-sheet conformation and disordered structures, with minimal α-helix content (<2%), essential for antibody’s biological functions (Table 1). The consistent CD spectra across variants suggest similar folding patterns of the R99 mAb regardless of the expression system employed.

**Table 1.**
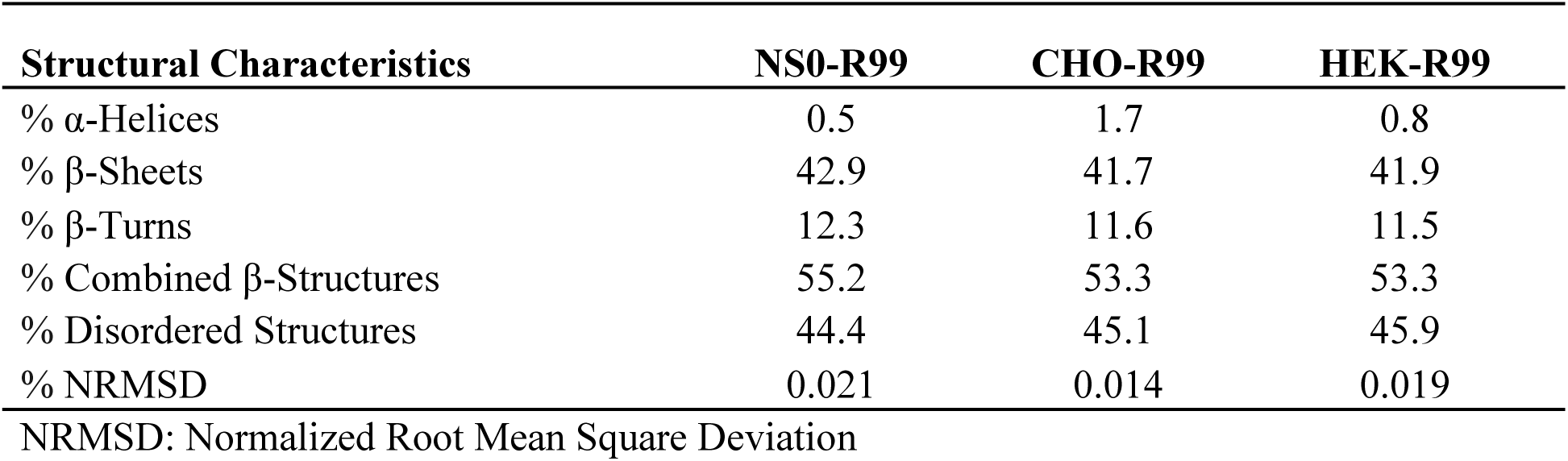
Secondary structure of R99 mAb derived from different host cell lines.

| **Structural Characteristics** | **NS0-R99** | **CHO-R99** | **HEK-R99** |
| --- | --- | --- | --- |
| % α-Helices | 0.5 | 1.7 | 0.8 |
| % β-Sheets | 42.9 | 41.7 | 41.9 |
| % β-Turns | 12.3 | 11.6 | 11.5 |
| % Combined β-Structures | 55.2 | 53.3 | 53.3 |
| % Disordered Structures | 44.4 | 45.1 | 45.9 |
| % NRMSD | 0.021 | 0.014 | 0.019 |

NRMSD: Normalized Root Mean Square Deviation

### R99 mAb showed a distinct charge profile and glycosylation pattern across host cell lines

Cation exchange chromatography revealed distinct charge profiles among the R99 variants (Table 2, Supplemental Figure 4). CHO-derived R99 exhibited the highest charge heterogeneity, with only 61.2% of species in the main peak, in sharp contrast with NS0-R99, characterized by a 91.9% distribution in the main peak and the lowest content in acid and basic species. HEK-derived R99 showed an intermediate charge profile with 80.8% distribution in the main peak. The high ionic strength required for R99 elution further indicated the presence of basic residues on the protein surface, consistent with its high content in arginine, closely associated with its specificity and biological function (Soto et al., 2012). Also, the treatment with carboxypeptidase B demostrated that the basic species observed in CHO-K1 and HEK-293 variants were not related to terminal lysine residues but likely attributable to PTMs (Not shown).

**Table 2.** Charge profile of R99 variants derived from different host cell lines.

| **Cation Exchange Parameters** | **NS0-R99** | **CHO-R99** | **HEK-R99** |
| --- | --- | --- | --- |
| % Main Peak | 93.9 | 61.2 | 78.7 |
| % Acidic Species | 5.5 | 21.4 | 16.5 |
| % Basic Species | 0.6 | 17.4 | 4.8 |

As expected, the glycosylation analysis highlighted significant differences among the variants (Table 3, Supplemental Figure 5). NS0-derived R99 exhibited a complex and highly glycosylated structure, with G1F (38.29%) and G2F (30.44%) as the predominant glycan form as compared to CHO and HEK variants. Additionally, NS0-R99 showed higher terminal galactose (83.07%) and sialylation (15.33%), indicating a more mature glycosylation pattern. In contrast, CHO-R99 and HEK-R99 had simpler glycosylation patterns, characterized by higher G0F content and lower levels of G1F, G2F, terminal galactose, and sialylation, while the fucosylation level was high across expression systems.

**Table 3.** Glycosylation profile of R99 mAb derived from different host cell lines.

| **Glycosylation Parameters** | **NS0-R99** | **CHO-R99** | **HEK-R99** |
| --- | --- | --- | --- |
| % G0F | 10.39 | 61.25 | 79.27 |
| % G1F | 38.29 | 19.81 | 12.95 |
| % G2F | 30.44 | 2.87 | 0.71 |
| % Fucosylation | 93.35 | 84.46 | 87.10 |
| % Terminal Galactose | 83.07 | 23.17 | 13.69 |
| % Sialylation | 15.33 | 0.59 | 0.02 |
| % High Mannose | 0.42 | 0.88 | 0.13 |

G0F: No glycosylated form, G1F: monogalactosylated form, G2F: digalactosylated form.

### The new recombinant variants of R99 mAb retained the specificity of the original mAb

Previous research has demonstrated that the R99 mAb binds with high affinity to its anti-idiotype mAb 1E10 (Fernández-Marrero et al., 2011) and recognizes sulfated GAGs, particularly CS, which is crucial for reducing subendothelial lipoprotein retention and atherosclerosis (Soto et al., 2012; Soto et al., 2024). To evaluate whether the new recombinant variants of R99 mAb preserve antigen recognition, we conducted an ELISA assessing their reactivity to both 1E10 mAb and CS. As depicted in Figure 2, both CHO-R99 and HEK-R99 mAbs showed comparable reactivity to 1E10 (Figure 2A) and CS (Figure 2B) relative to the parental NS0-R99. The observed dose-dependent recognition further indicates that the new variants retain their antigen specificity across a broad range of concentrations. As expected, the negative control hR3 mAb showed no reactivity, highlighting the specificity of the new variants. These findings confirm that both CHO-R99 and HEK-R99 mAbs preserve the essential antigen-binding properties of the original NS0-R99 mAb, suggesting their potential for therapeutic applications targeting GAGs and related antigens. Given the critical role of sulfated GAGs in atherosclerosis, we investigated whether the new R99 variants could recognize atherosclerotic lesions from NZW rabbits. As illustrated in Figure 2C, the mAbs derived from different cell lines exhibited a similar pattern of recognition in consecutive sections of rabbit aortas. The mAbs specifically targeted atherosclerotic lesions, while no reactivity was observed in aortic sections exposed to the hR3 isotype control mAb.

**Figure 2.**
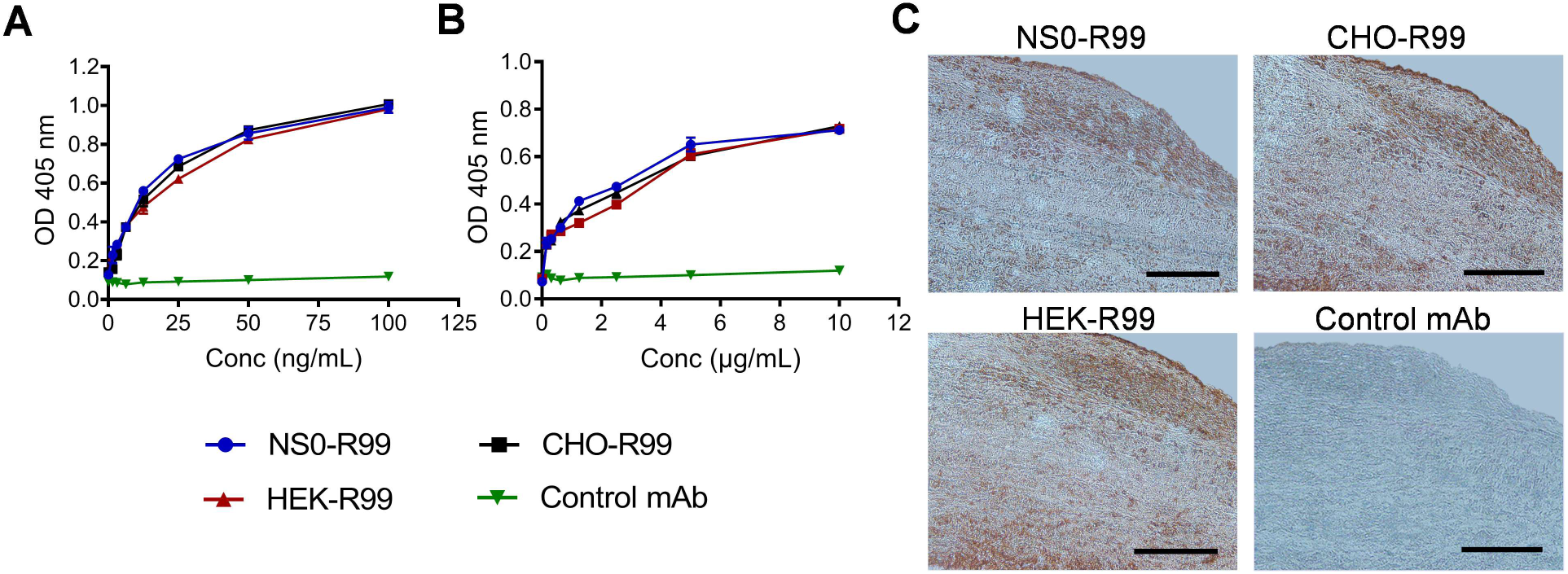
Recognition properties of the R99 mAb produced in CHO-K1 and HEK293 cell lines. (A) Reactivity of R99 to 1E10 mAb by ELISA, as example of anti-idiotypic (Ab2) mAb of R99. 1E10-coated plates were incubated with the mAbs (1.5-100 ng/mL) followed by an alkaline phosphatase-conjugated anti-human IgG secondary antibody. (B) Comparative recognition of R99 variants to chondroitin sulfate by ELISA. Chondroitin sulfate-coated plates (10 µg/mL) were incubated with different concentrations of the mAbs (0.15-10 µg/mL) followed by an alkaline phosphatase-conjugated anti-human IgG secondary antibody. (C) Recognition of R99 variants to rabbit’s aorta. Consecutive aortic sections from NZW rabbits treated with Lipofundin 20% were incubated with 20 µg/mL of the R99 mAb. Binding was detected using a goat anti-human IgG antiserum conjugated to biotin followed by a horseradish peroxidase-streptavidin complex. Magnification 20x, Bar = 100 µm.

### R99 new variants preferentially accumulated in rat aorta thereby reducing LDL retention

The efficacy of R99 mAb variants in reducing LDL retention and oxidation was assessed in rats *in vivo* (Calara et al., 1998; Soto et al., 2012). For this study, we administered 1 mg of the mAbs (approximately 4 mg/kg, equivalent to 270 mg in humans); a dose comparable to those typically employed in clinical passive mAb therapies (Figure 3A). Additionally, we evaluated the biodistribution of R99 in healthy animals at this high dose to determine specificity and rule out potential off-target effects under normal physiological conditions. Interestingly, serum quantification of human IgG showed that circulating levels of both R99 mAb variants were below 10 µg/mL, 4-fold lower than the one observed for the isotype control mAb hR3(Figure 3B). The faster clearance of R99 mAb suggested its preferential accumulation in specific tissues. Indeed, quantitative ELISA in organ homogenates revealed that both CHO-R99 and HEK-R99 mAbs accumulated in normal aorta, considering specific accumulation when R99 levels were at least 2-fold higher than the isotype-matched mAb (Figure 3C). Importantly, hR3 was chosen as a suitable isotype-matched control due to its specificity for the human EGF receptor and its lack of cross-reactivity with other species, providing a baseline for human IgG clearance and organ accumulation in rats. To further assess the impact of R99 arterial accumulation on LDL retention and oxidation, we performed immunohistochemical analysis using the EO6 mAb on aortas from rats administered LDL (Figure 3D-E). Oxidized LDL was detected throughout the arterial wall in rats treated with PBS or hR3 mAb. In contrast, the R99 new variants significantly reduced LDL retention and oxidation, consistent with previous findings for the NS0 parental mAb (Soto et al., 2012).

**Figure 3.**
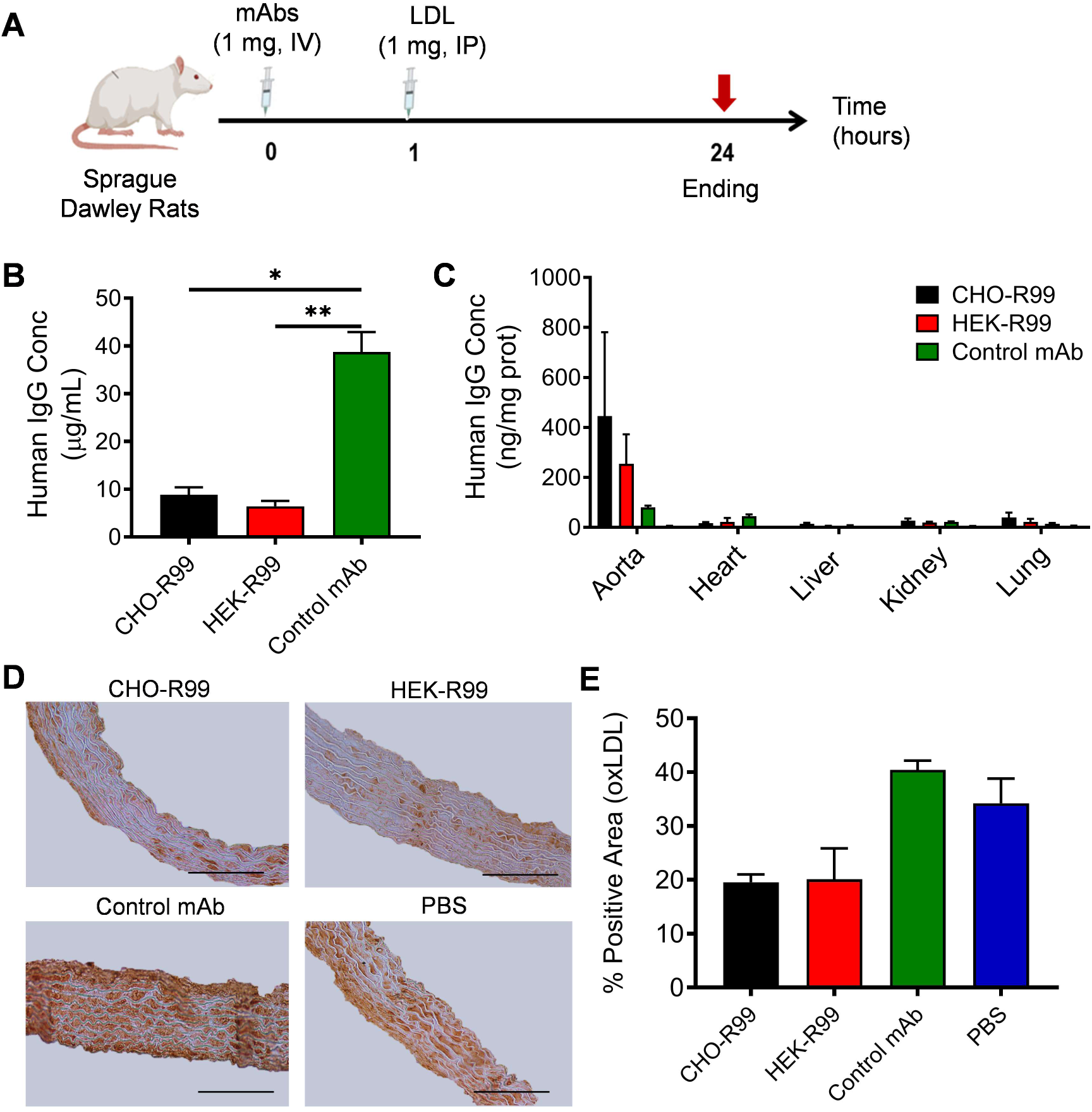
Blocking properties of the R99 mAb produced in CHO-K1 and HEK293 cell lines on LDL arterial retention/oxidation *in vivo*. (A) Sprague Dowley rats (n=3) received a 1 mg IV injection of CHO-R99, HEK-R99, or hR3 as a human control mAb, as well as 1 mL of PBS in a control group. After one hour, all groups were administered LDL (1 mg, IP). Quantification of the mAbs in (B) sera and (C) tissue lysates was performed by ELISA, revealing a fast clearance of R99 variants from the circulation and their specific accumulation in aorta. (D) Representative images of the blocking properties of the R99 variants on LDL retention/oxidation in rats. The presence of oxidized LDL in the arterial wall was determined by incubating aortic sections with the commercial antibody EO6. Reactivity was detected with a biotin-conjugated goat anti-mouse immunoglobulin polyclonal antiserum (1:200; Jackson) followed by horseradish perosidase-conjugated streptavidin complex (1:200; Jackson). Magnification 20X. Bars=100 µm. (E) Percentages of EO6 mAb positive area relative to the total tissue area. Data is represented as the mean±SEM, *p<0.05, **p<0.01, Kruskal-Wallis’ test followed by Dunn’s multiple comparison.

### Passive administration of recombinant R99 mAbs reduced atherosclerosis in NZW rabbits

#### R99 variants accumulate in aorta in a NZW rabbit model of atherosclerosis

The biodistribution of CHO-R99 and HEK-R99 mAbs was evaluated *in vivo* under proatherogenic conditions in NZW rabbits undergoing Lipofundin IV administration. Over a 10-day period, rabbits received a total of 6 mg of mAbs, and we quantified human IgG levels in blood and organ homogenates (Figure 4A). As shown in Figure 4B, R99 variant levels in circulation were negligible by the end of the study, indicating significant tissue accumulation and rapid clearance from the bloodstream. In contrast, the isotype control mAb hR3 maintained a serum concentration of 25.8 ± 4.1 µg/mL, reflecting a slower clearance compared to the R99 variants. Biodistribution analysis in rabbits (Figure 4C) revealed that both CHO-R99 and HEK-R99 mAbs displayed similar accumulation patterns across various organs. Both variants preferentially accumulated in the aorta of rabbits treated with Lipofundin, with CHO-R99 and HEK-R99 levels at 10.7 ± 5.6 ng/mg and 11.3 ± 6.8 ng/mg of protein, respectively. This accumulation in the aorta was approximately 10-fold higher than in other organs, where mAb levels were very low. Examination of organs with high blood supply and those involved in mAb catabolism, including the heart, liver, kidneys, and lungs, confirmed the targeted accumulation of mAbs in the aorta.

**Figure 4.**
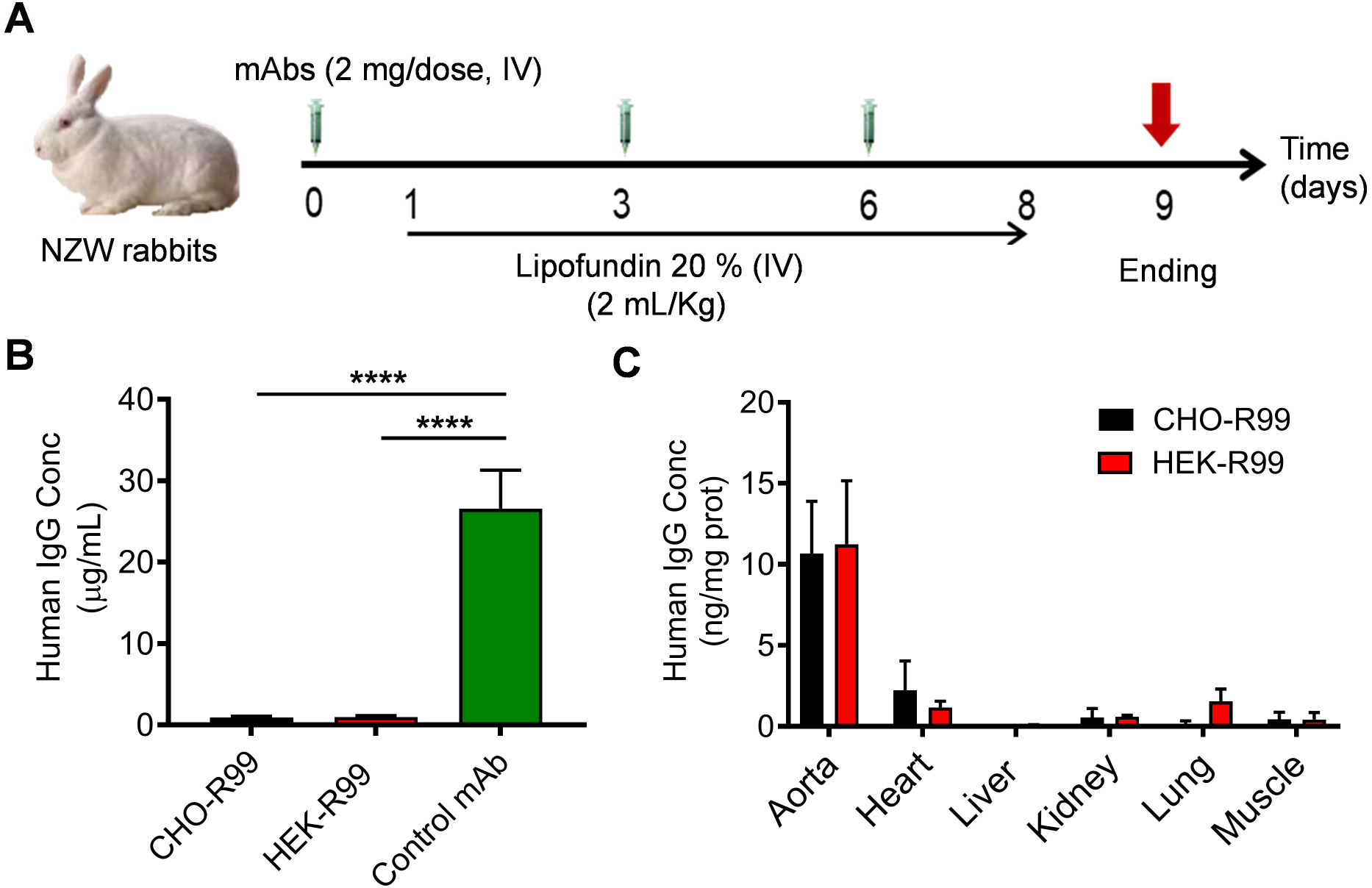
Biodistribution of the R99 mAb produced in CHO-K1 and HEK293 cell lines in NZW female rabbits treated with Lipofundin 20%. (A) The animals (n=6-7 per group) were given three IV injections of the mAbs in PBS every three days (2 mg/dose), concurrently with the induction of atherosclerosis by a daily IV administration of Lipofundin 20% (2 mL/kg, 8 days). Quantification of the mAbs in (B) sera and (C) tissue lysates was performed by ELISA, revealing a fast clearance of R99 variants from the circulation and their specific accumulation in aorta. Data is represented as the mean±SEM, ****p< 0.0001, one-way ANOVA followed by Tukey’s multiple comparisons test.

#### R99 variants exerted an antiatherogenic effect in rabbits

The antiatherogenic effect of the new R99 mAb variants was assessed in an acute model of atherosclerosis developed in NZW rabbits by a daily administration of the lipid emulsion Lipofundin 20% (Delgado Roche et al., 2012; Medina et al., 2012). Each rabbit received a total mass of 6 mg of R99 mAbs troughout a 10 days intervention period, equivalent to 68 mg in humans. The treatment with either mAb did not caused apparent abnormalities in major organs. Histological analysis of the aortas on H&E and Alcian Blue stainings (Figure 5A) revealed that rabbits receiving the isotype control mAb hR3 developed significant intimal thickening, irregular luminal surfaces, marked lipid accumulation, and smooth muscle cell migration into the intima, along with increased GAG content and extracellular matrix deposition. In contrast, rabbits treated with CHO-R99 showed intimal thickening in two animals and no significant changes in the remaining four. The HEK-R99 group also exhibited reduced lesion formation, with fewer lesions than the control in the three out of seven rabbits treated. Specifically, the GAG content in both R99-treated groups was lower compared to the control mAb. Lesions were categorized as mild, moderate, or advanced based on intimal thickening assessed by Verhoeff-Van Gieson staining (Suresh et al., 2020) along with the extracellular matrix content and the presence of extracellular lipids. Approximately 60% of rabbits treated with either R99 variant did not develop atherosclerosis, compared to 100% in the isotype control group. No moderate or advanced lesions were observed in the CHO-R99 group, while 29% of the HEK-R99 group developed moderate lesions. In contrast, 83% of the control group exhibited moderate to advanced lesions. Quantitative analysis of intimal thickening revealed that the R99-treated groups had significantly fewer and smaller lesions, with 7 and 10 significant lesions in the CHO-R99 and HEK-R99 groups, respectively (p<0.05), compared to 22 lesions in the control group (Figure 5D).

**Figure 5.**
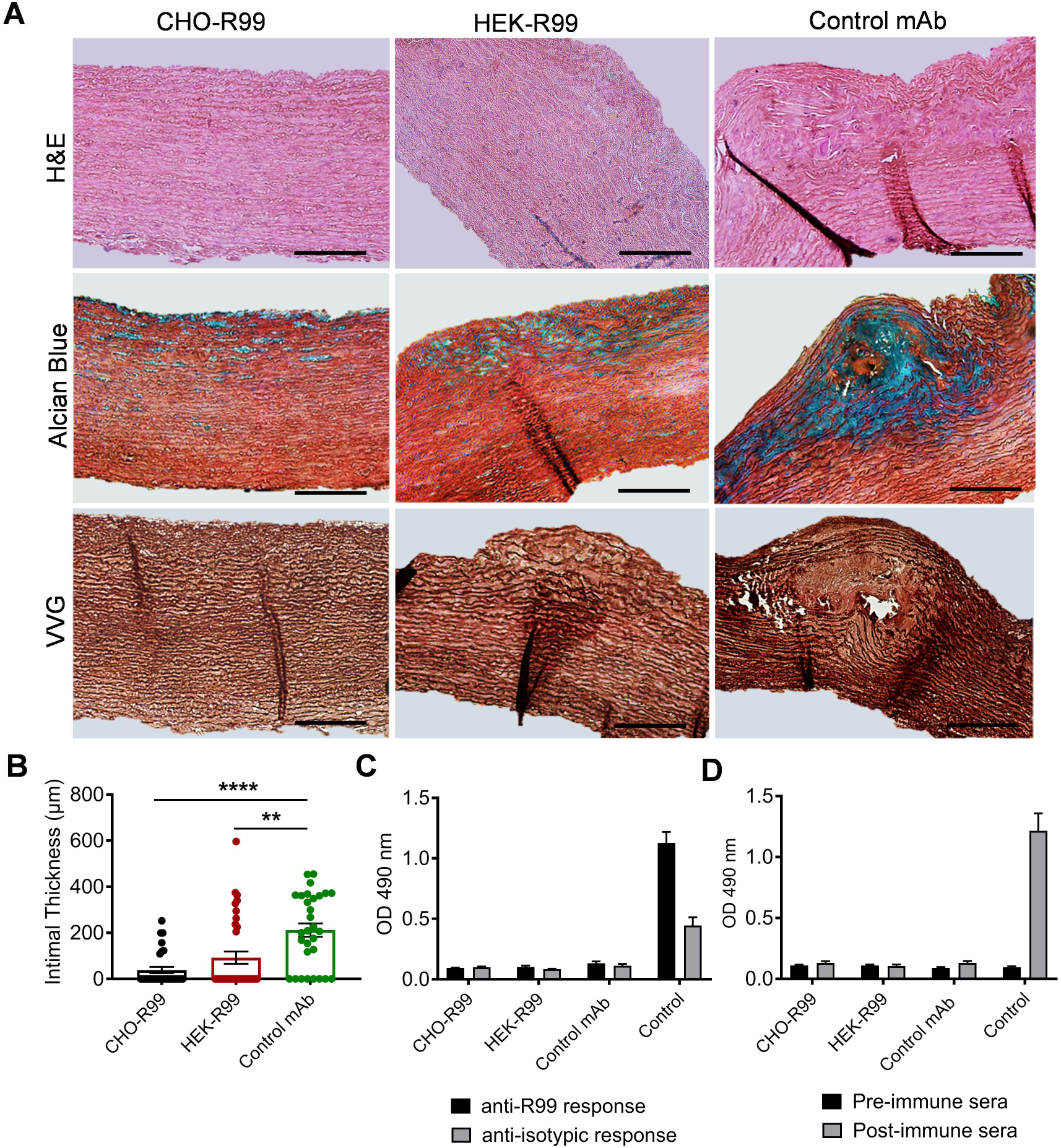
Antiatherogenicity of R99 mAb produced in CHO-K1 and HEK293 cell lines on Lipofundin-induced atherosclerosis in female NZW rabbits. The animals (n=6-7 per group) were given three IV injections of the mAbs in PBS every three days (2 mg/dose), concurrently with the induction of atherosclerosis by a daily IV administration of Lipofundin 20% (2 mL/kg, 8 days). (A) Histopathological analysis from H&E, Alcian Blue and Verhoeff-Van Giesson-stained aortic sections revealed intimal thickening and accumulation of glycosaminoglycans (blue) in the isotype control-treated group. Both R99 variants reduced the lesion size and the extracellular matrix secretion. (B) Intimal thickness from five different aortic regions per animal. (C) Anti-idiotype (Ab2) IgG response in NZW rabbits. ELISA plates coated with 10 µg/mL of R99 or hR3 mAbs were incubated with the post-immune sera (dil 1/5000). (D) Anti-chondroitin sulfate (Ab3) IgG response in NZW rabbits. Chondroitin sulfate-coated plates (10 µg/mL) were incubated with the pre-inmune and post-immune sera (1/400) followed by a horseradish peroxidase-conjugated anti-rabbit IgG secondary antibody. Sera from seven rabbits previously immunized with NS0-R99 mAb were used as immunogenicity controls. Data are mean±SEM. **p<0.01, ****p<0.0001, Kruskal-Wallis’ test followed by Dunn’s multiple comparison test.

#### The antiatherogenicity of the R99 variants in rabbits was not associated with the induction of an anti-idiotypic cascade

The induction of anti-idiotype (Ab2) and anti-chondroitin sulfate (Ab3) antibodies in NZW rabbits was evaluated by ELISA. Figure 5E shows that no Ab2 antibodies were detected against either the isotype control (grey bars) or the idiotype of the R99 mAb (black bars). Similarly, Figure 5F indicates that there was no generation of Ab3 (anti-CS antibodies). This indicates that, over the 10-day period, there was no significant antibody cascade triggered by the intravenous administration of the R99 mAb. In contrast, control rabbits immunized with the R99-NS0 mAb using a regimen of four weekly SC injections used as a control, exhibited detectable Ab2 and Ab3 responses.

## DISCUSSION

According to the “Response to Retention” hypothesis proposed by Williams and Tabas in 1995, the retention of LDL in the subendothelial space, mediated by proteoglycans, is a critical event in the onset and progression of atherosclerosis (Williams and Tabas 1995; Camejo et al., 1998; Skålén et al., 2002; Tabas et al., 2007). It is now well established that, in addition to LDL, other ApoB-containing lipoproteins, such as lipoprotein (a) and triglyceride-rich lipoproteins— including dietary-derived chylomicron remnants—also play significant roles in atherogenesis (Williams and Tabas 1995; Williams and Fisher 2015). Despite the relevance of these mechanisms, therapies targeting the vascular wall components involved in lipoprotein retention remain largely underexplored (Zeng et al., 2005; Ballinger et al., 2010; Kumarapperuma et al., 2024). To address this, we have developed the chimeric R99 mAb, which recognizes sulfated GAGs and targets the arterial retention of lipoproteins (Soto et al., 2012). This mAb has demonstrated antiatherosclerotic effects in various experimental models, attributed to its ability to inhibit subendothelial LDL retention, either directly or through the induction of autologous antibodies with similar specificity (Brito et al., 2012; Soto et al., 2012; Delgado-Roche et al., 2015; Brito et al., 2017; Sarduy et al., 2017; Soto et al., 2024). Its dual functionality—targeting CSPGs and eliciting an antibody response—supports both passive therapy and vaccination approaches (Soto et al., 2024). While R99 mAb effectively blocks arterial lipoprotein retention *in vivo*, current studies in animal models have primarily focused on idiotypic vaccination with low-dose SC administrations. Passive therapies require high doses of mAbs over extended periods, necessitating efficient strategies for recombinant cell line development in serum-free media to meet production demands (Butler 2015; Kelley 2020).

Although R99 mAb is currently expressed in NS0, CHO-K1, and HEK-293 cell lines, a detailed analysis of their structural characteristics in relation to their efficacy against atherosclerosis has not been previously conducted. In this study we performed a comprehensive comparative physicochemical characterization of the R99 variants to investigate how expression systems, particularly PTMs, might affect the antibody’s structural integrity, antigen-binding specificity, and overall functional performance. Given that R99 mAb’s complementarity-determining regions (CDRs) are rich in arginines, which are critical for antigen recognition and antiatherogenic activity, our study specifically examined whether expression-related differences could influence its efficacy in atherosclerosis models.

### Comparative structural characteristics of R99 mAb expressed in different cell lines

Physicochemical characterization of R99 mAb variants expressed in NS0, CHO-K1, and HEK-293 cells demonstrated consistent structural integrity and functional stability across all expression systems. All variants displayed the expected molecular weights of 150 kDa for the intact mAb, 25 kDa for the light chain, and 50 kDa for the heavy chain. Additional bands observed in electrophoretic profiles suggested the presence of isoforms and/or PTMs, which are common in antibody production (Liu et al., 2019). Despite these variations, SEC-HPLC indicated high purity, with over 99% monomeric fractions and minimal aggregation in all variants, consistent with established standards (Kirley and Norman 2018). DLS further confirmed monodisperse preparations with good colloidal stability, a critical attribute for therapeutic antibodies.

Primary and secondary structure analyses revealed subtle PTM differences among variants. Peptide mapping indicated largely conserved sequences across the different expression systems, with NS0-derived mAb showing additional peaks, potentially due to differential PTMs such as asparagine deamidation, aspartic acid isomerization, methionine and tryptophan oxidation, lysine glycation, and glycosylation across variants (Li et al., 2015; Xu et al., 2019). The analysis was performed using a combination of trypsin and Lys-C enzymes, which cleave at arginine and lysine residues, respectively. Given the high prevalence of arginine residues within the CDRs of R99 mAb, future studies could benefit from alternative enzymes like Glu-C or Asp-N to preserve key antigen-recognition regions. Nevertheless, far UV CD spectroscopy confirmed consistent secondary structures across all variants, with similar β-sheet and disordered structures percentages, suggesting that the core folding and structural integrity, essential for antigen-binding, were preserved (Sissolak et al., 2019). This indicates that the observed primary structure variations may have minimal impact on stability and recognition properties, aligning with reports that such modifications often do not significantly alter the therapeutic performance of antibodies.

Cation exchange chromatography further highlighted differences in charge profiles among the R99 variants, primarily due to PTMs as suggested by peptide mapping. The NS0-derived variant displayed a relatively uniform charge profile, with a main peak representing 93.4% of total species, suggesting fewer PTMs and reduced isoform formation. The CHO-K1 variant, however, exhibited greater charge heterogeneity, with the main peak accounting only for only 61.2% of total species, reflecting a higher presence of both acidic and basic isoforms, likely due to glycosylation variations and PTMs like deamidation or isomerization (Kabat 1991). The HEK-293-derived variant showed an intermediate charge profile, with the main peak constituting 78.7% of total species. Additional, carboxypeptidase B treatment, which removes terminal lysine residues, did not significantly alter charge profiles, suggesting that basic species are not primarily due to these residues but rather other PTMs such as N-terminal pyroglutamate formation or asparagine deamidation (Hermanson 2013). This observation is consistent with literature, which reports that endogenous carboxypeptidases in cell culture media efficiently cleave these residues, thereby minimally impacting charge heterogeneity (Zhang et al. 2015). The overall high basic charge content identified on the surface of R99 mAb, as determined by the high ionic strength required to elute the peptides, is also consistent with the abundance of arginine residues in the CDRs, reported for the mAb’s affinity to sulfated GAGs. While charge heterogeneity did not significantly impact antigen-binding (Figure 2), the presence of multiple isoforms could pose challenges for product consistency, affecting solubility, aggregation, and pharmacokinetics (Sissolak et al., 2019).

In line with peptide mapping and charge profile analyses, glycosylation patterns accounted for the primary differences among R99 variants, consistent with literature showing that glycosylation profiles are heavily influenced by the expression system, with additional modulation by cell culture conditions (Walsh and Jefferis 2006; Jenkins et al., 2008; Hossler et al., 2009) (Hossler et al., 2009; Jenkins et al., 2017; Walsh and Jefferis, 2006). NS0-derived R99 exhibited a complex glycosylation pattern with high levels of terminal galactose, sialylation, and a greater proportion of G1F and G2F glycans, reflecting the advanced glycan processing capacity of NS0 cells. These characteristics could not only enhance undesired effector functions in the context of atherosclerosis, such as antibody-dependent cellular cytotoxicity and complement-dependent cytotoxicity (Jenkins et al., 2008), but also introduce additional safety concerns. These include non-human glycan epitopes, such as N-glycolylneuraminic acid and α-galactose (Galα(1,3)Galβ(1,4)GlcNAc), which have been linked to adverse immune reactions (Ghaderi et al., 2010). In contrast, CHO-K1 and HEK-293 variants showed simpler and more similar glycosylation profiles, with core-fucosylated, biantennary structures that are less complex and more human-like in HEK-293, potentially reducing the risk of adverse immune reactions and offering advantages in pharmacokinetics (Walsh and Jefferis 2006; Hossler et al., 2009). However, the immunogenicity of R99 mAb is not solely determined by its glycosylation and other PTMs, but also by the well-documented immunodominance of its idiotype, which is central to its mechanism of action and therapeutic effect (Brito et al., 2012; Sarduy et al., 2017). Therefore, future studies are essential to assess the immunogenicity of R99 across variants to fully understand the impact of glycan differences on efficacy and safety. Additionally, the simplicity of CHO-K1 and HEK-293 glycosylation patterns could also impact solubility and stability, which could be a concern for maintaining product consistency across batches.

Despite these structural differences, ELISA and immunohistochemical assays demonstrated that all R99 variants maintained comparable antigen recognition and relative binding affinity to anti-idiotypic antibodies, sulfated GAGs, and rabbit atherosclerotic lesions (Figure 2), consistent with previous studies on NS0-R99 mAb (Fernández-Marrero et al., 2011; Soto et al., 2012; Soto et al., 2014). These findings suggest that the CDRs, critical for antigen recognition, remained largely unaffected by PTM variations, underscoring the robustness and consistent functional performance of R99 mAb.

### Biodistribution and antiatherogenic properties of the R99 variants expressed in CHO-K1 and HEK-293 cells

After confirming the structural integrity and preserved reactivity of the CHO-R99 and HEK-R99 variants for their target antigen, we evaluated their biological efficacy and antiatherogenic potential *in vivo*. First, we conducted biodistribution studies in both healthy rats and a rabbit model of experimentally induced atherosclerosis. This dual approach was designed to consider the expression of CSPGs in various tissues under normal physiological conditions, their overexpression during atherosclerosis, and the limited safety data associated with specific treatments targeting these molecules (Wight, 2018). In both species, the CHO-R99 and HEK-R99 mAbs demonstrated consistent and highly specific accumulation in arterial tissues, with minimal off-target deposition. Notably, the rapid clearance of R99 variants from circulation, coupled with their preferential accumulation in arterial tissues, reinforce their potential for targeting atherosclerotic plaques without systemic toxicity. This specificity was further confirmed by comparisons with the isotype-matched control hR3, which does not cross-react with non-human species and served as a reference for human IgG clearance and organ distribution. It is worth noting that up to date, no toxic effect has been observed in any of the species studied, including mice, rats, and rabbits (Brito et al., 2012; Soto et al., 2012; Sarduy et al., 2017; Soto et al., 2024). Our studies in healthy Sprague Dawley rats are consistent with recent findings demonstrating that IV administration of NS0-R99 at a dose of 4.5 mg/kg did not adversely affect lipid metabolism (both fasting and postprandial), glucose and insulin resistance indices, liver or kidney function, blood cell counts, or systemic levels of cytokines and growth factors in both insulin-resistant and wild-type rats (Soto et al., 2024). The observed biodistribution pattern in rabbits also aligns with previous studies involving NS0-R99 mAb, which demonstrated specific uptake in atherosclerotic lesions in rabbits and mice (Soto et al., 2014; Brito et al., 2017).

These results reinforce the specificity of R99 mAb for arterial CSPGs, particularly due to the structural characteristics of CSPGs under pathological conditions (Little et al., 2007). In atherosclerosis, CSPGs are mostly synthesized by vascular smooth muscle cells, displaying notable structural modifications, such as elongated GAG chains and altered sulfation patterns, which increase their affinity for LDL (Ballinger et al., 2007; Little et al., 2007). The enhanced binding of the R99 mAb variants to these modified CSPGs in atherosclerotic lesions likely contributes to their targeted accumulation in arterial tissues, while CSPGs in other tissues are produced by distinct cell types and possess different structural and functional characteristics (Kwon et al., 2008, Wight, 2018). Their consistent structural pattern across vertebrates further emphasizes the translational relevance of using different animal models to target arterial CSPGs with R99 mAb when addressing safety and efficacy on the treatment for atherosclerosis (Yamada et al., 2011).

The efficacy of the R99 mAb variants was evaluated in parallel using the lipoprotein retention model in rats and the acute atherosclerosis model in rabbits, alongside the biodistribution studies conducted for these animals. The dose selection in each model was guided by allometric scaling principles to ensure translational relevance to human dosing (Nair et al., 2016; Germovsek et al., 2021).

In the rat model, we administered the mAbs at an approximate dose of 4 mg/kg, corresponding to a human equivalent dose of about 270 mg. This high dose was chosen to robustly evaluate the ability of the R99 variants to block LDL retention in the arterial wall, a critical mechanism of atheroprotection previously described for NS0-R99 (Soto et al., 2012). This dosing strategy aligns with typical clinical mAb therapies, which generally range from 75 mg to 300 mg per administration, with higher doses or increased frequency often required to achieve sufficient therapeutic efficacy in passive immunotherapy (Gluba-Brzózka et al., 2021; Dankar et al., 2024). Equivalent protein masses of LDL and R99 were used to assess their competitive binding over 24 h under similar conditions. The efficacy of the R99 variants in reducing LDL retention/oxidation in the arterial wall was confirmed by immunohistochemical analysis using the EO6 mAb, which specifically recognizes oxidized phospholipids. In control rats treated with PBS or the hR3 mAb, oxidized LDL was extensively detected throughout the arterial wall. However, administration of the CHO-R99 and HEK-R99 variants resulted in a reduction in LDL retention and oxidation, achieving an approximate 50% decrease compared to the control groups. This effect is consistent with previous findings for the NS0-R99 mAb in Sprague Dawley rats (Soto et al., 2012), indicating that the R99 variants retain comparable affinity for arterial CSPGs and effectively reduce LDL deposition and oxidation in the arterial environment. These findings also align with previous observations that NS0-R99 demonstrated a strong competitive effect in solid-phase assays, blocking approximately 50% of LDL and chylomicron remnant binding to CS and vascular extracellular matrix *in vitro*, when equivalent particle numbers of lipoproteins and R99 mAb were tested. Moreover, pre-perfusion with NS0-R99 mAb in insulin-resistant rats resulted in specific antibody accumulation within the arterial wall and a dose-dependent reduction in fluorescent lipoprotein retention, achieving around a 60% decrease in lipoprotein binding and cholesterol deposition under similar conditions (Soto et al., 2024).

In contrast to the rat experiments, the rabbit model employed a lower dose of 3 mg/kg, equivalent to an estimated human dose of 68 mg. This dose was intentionally lower than typical clinical doses for mAbs in cardiovascular diseases, which often range from 120-150 mg administered biweekly or monthly, leading to cumulative doses of 300-400 mg (Dixon et al., 2019). The rationale for selecting a lower dose in rabbits was to challenge the potential antiatherogenic effects of the R99 variants in an acute atherosclerosis setting, allowing us to discern differences in efficacy between the CHO-R99 and HEK-R99 variants.

In our rabbit studies, Lipofundin 20% has been used as an atherosclerosis-inducing agent, based on IV administration of this lipid emulsion over eight days to induce acute atherosclerotic changes and systemic and arterial oxidative stress (Noa and Más 1992; Delgado Roche et al., 2012; Medina et al., 2012). This study showed that the R99 mAb variants produced in both CHO-K1 and HEK-293 cell lines effectively recognize GAGs and accumulate in atherosclerotic lesions *in vivo*, supporting their potential use for passive atherosclerosis treatment. In control rabbits treated with the unrelated mAb hR3, Lipofundin 20% induced lesions characterized by intimal thickening, high GAG content, distorted smooth muscle cell arrangement, and extracellular lipid accumulation, consistent with our prior reports (Noa and Más 1992; Medina et al., 2012).

To evaluate the antiatherogenic efficacy of the R99 mAb, we performed morphometric analyses focused on the five largest intimal thickness measurements per animal, due to the focal and variable nature of atherosclerotic lesions. Both R99 variants significantly reduced intimal thickness and provided complete protection in 60% of the animals from developing atherosclerosis. While the total number of lesions was comparable between the variants, in the rabbits that developed atherosclerotic lesions, the group treated with CHO-R99 tended to exhibit less intimal thickening than the one receiving HEK-R99 mAb, suggesting potential differences in efficacy. Importantly, even at doses approximately half of those commonly used in clinical settings for atherosclerosis treatment, R99 mAbs demonstrated substantial protective effects, emphasizing their therapeutic potential. Although our results support the development of either CHO-R99 or HEK-R99 for passive treatment of atherosclerosis, minor differences in the number and lesion size between the variants indicate that further dose-response studies are required to determine the optimal therapeutic dose for passive immunotherapy and to maximize clinical outcomes.

Previous research has shown that the R99 mAb is highly immunogenic, with its immunodominant idiotype capable of inducing a cascade of anti-idiotype (Ab2) and anti-anti-idiotype (Ab3) antibodies, where Ab3 shares the specificity of R99 mAb for recognizing sulfated glycosaminoglycans (GAGs). This immunogenic property has been observed across various species and has contributed to the atheroprotective effects seen in immunization studies (Brito et al., 2012; Soto et al., 2012; Sarduy et al., 2017; Soto et al., 2024). Low, SC doses of R99 mAb have been demonstrated to induce a humoral response of Ab3 antibodies in a dose-dependent manner, independent of age or sex (Sarduy et al., 2017). Typically, the induction of Ab2 requires at least two weeks, with Ab3 appearing after three weeks, reaching a maximum response by the fourth week (Soto et al., 2024). For instance, in rabbit models, administering 100 µg of R99 mAb weekly over 3-4 weeks is necessary to observe these effects (Soto et al., 2012). However, the acute NZW rabbit model used in the referenced study generates lesions within eight days, a timeframe insufficient to mount an idiotypic cascade. Consequently, the absence of Ab2 and Ab3 antibodies confirmed that indeed the observed antiatherogenic effect in this context is attributable to the direct blocking action of the mAb rather than its immunogenic properties.

### Final considerations

In conclusion, this study provides the first evidence of the antiatherogenic effects of R99 mAb through passive administration in animal models. Despite variations in post-translational modifications among NS0, CHO-K1, and HEK-293 expression systems, all mAb variants exhibited consistent antigen recognition, specificity for arterial tissues, and therapeutic efficacy *in vivo*. Notably, even at lower doses, the R99 variants significantly reduced atherosclerotic lesion development, underscoring their potential for passive immunotherapy in cardiovascular diseases. While the NS0 variant’s more complex glycosylation might increase immunogenicity, potentially enhancing its use as a vaccine, the CHO-K1 and HEK-293 variants offer scalable manufacturing with favorable safety profiles for passive therapies. These findings validate the therapeutic potential of chP3R99 mAbs and emphasize the need for further immunogenicity and dose-response studies to optimize their clinical application.

## ACKNOWLEDGEMENTS

We are grateful for the technical expertise provided by Ms. Tania Gómez.

## SUPPLEMENTAL MATERIAL

### Supplemental Figures

**Supplemental Figure 1.**
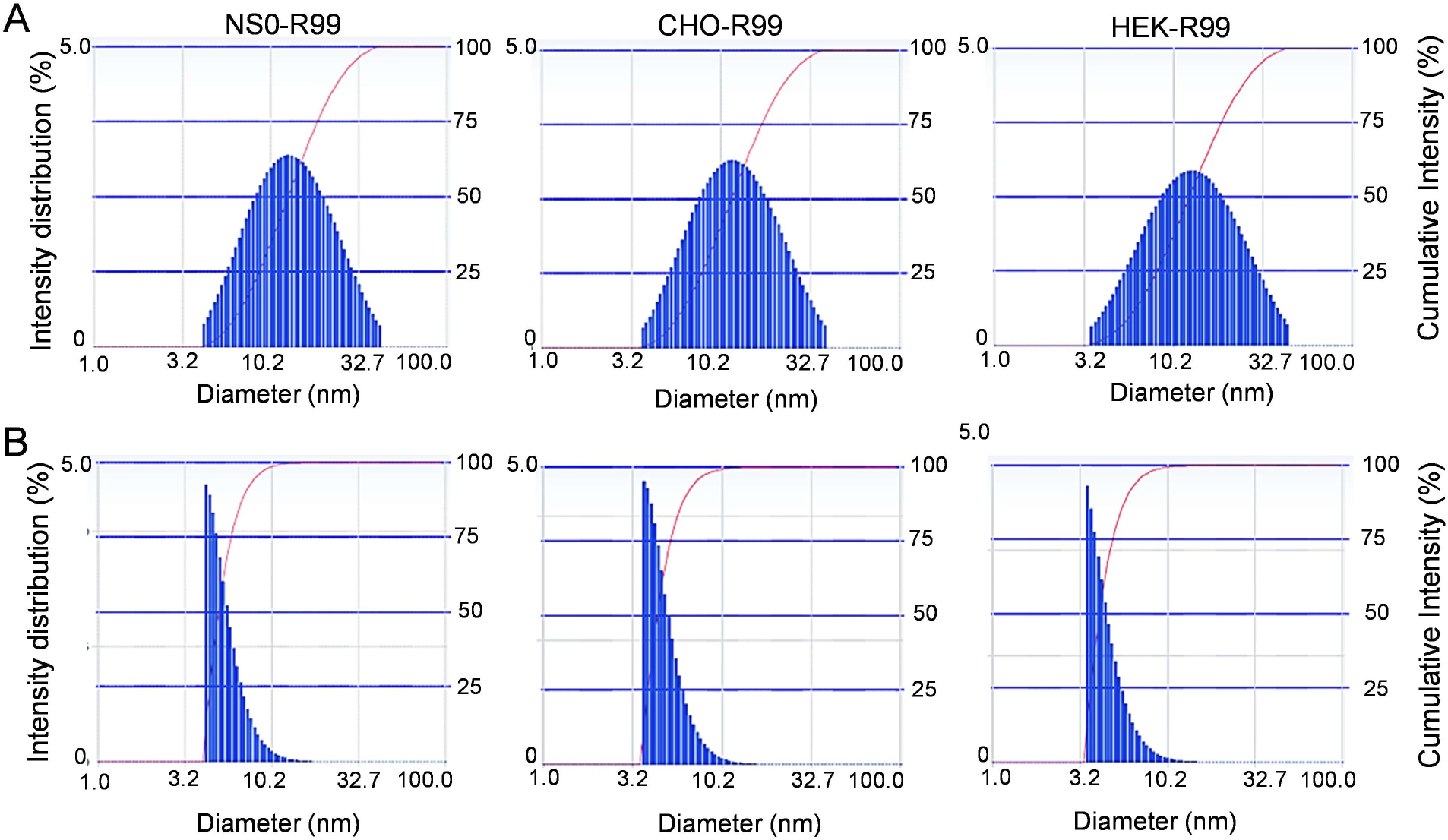
Particle size measured by Dynamic Light Scattering of R99 mAb expressed in NS0, CHO-K1, and HEK-293 host cell lines. (A) Intensity distribution graphs and (B) number distribution graphs. Data acquisition involved triplicate measurements of 100 counts each, utilizing Cumulant and Contin statistical methods.

**Supplemental Figure 2.**
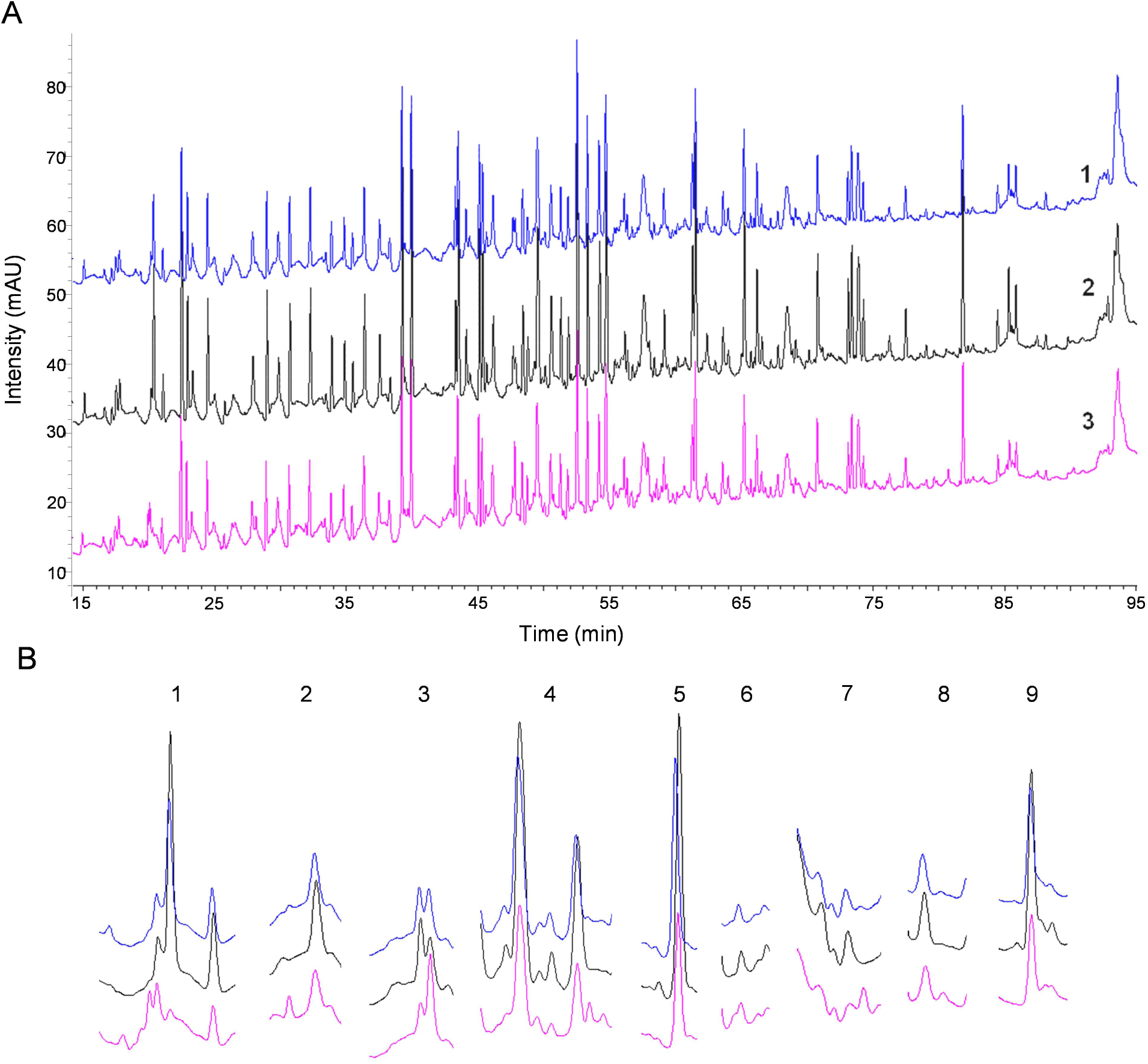
Representative peptide mapping profiles of R99 mAb expressed in NS0, CHO-K1, and HEK-293 host cell lines obtained by RP-HPLC following trypsin/Lys-C digestion. A) The chromatograms correspond to different expression systems: NS0-R99 (magenta), CHO-R99 HEK (blue), and HEK-R99 NS0 (black). Each profile reflects the specific peptide fragments generated, allowing for comparison of the mAb variants across expression systems. B) Main regions of the chromatogram where differences were identified. Two batches of each variant were analyzed.

**Supplemental Figure 3.**
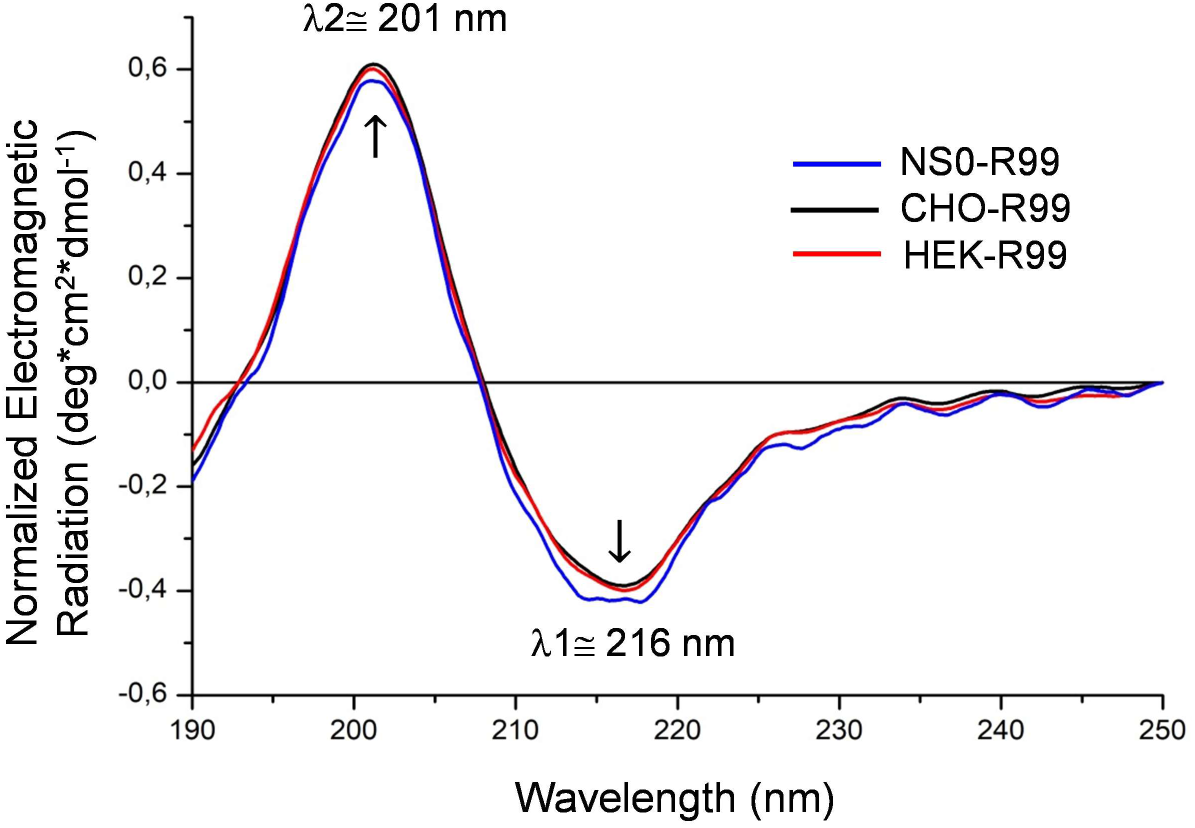
Representative Far-UV circular dichroism (CD) spectra of R99 mAb expressed in NS0, CHO-K1, and HEK-293 host cell lines. The spectra show the mean residue ellipticity (deg·cm²·dmol⁻¹) as a function of wavelength at 25°C for each variant: NS0-R99 mAb (blue), CHO-R99 mAb (black), and HEK-R99 mAb (red). Arrows indicate the characteristic positive band around 201 nm and the negative band around 216 nm, consistent with the β-sheet-rich secondary structure typical of these mAbs. Two batches of each variant were analyzed.

**Supplemental Figure 4.**
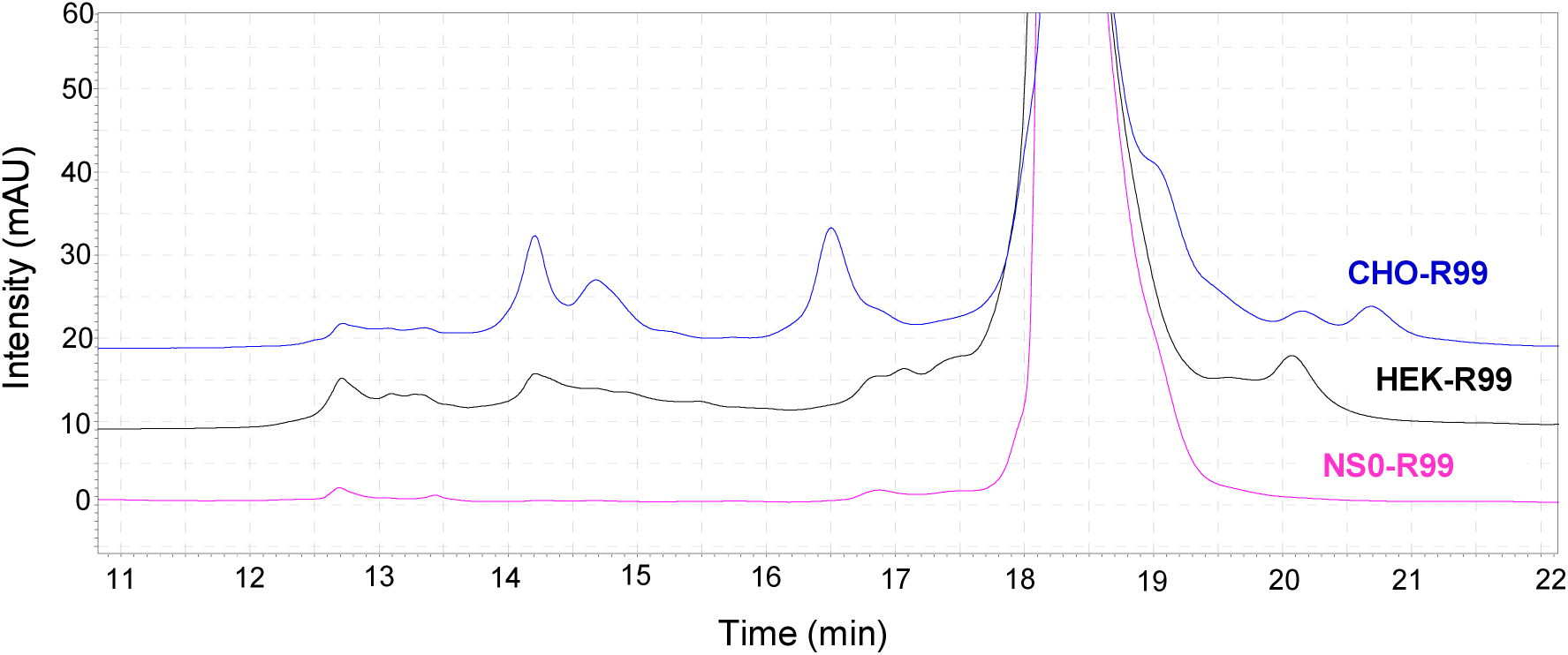
Representative charge profile of R99 mAb expressed in NS0, CHO-K1, and HEK-293 host cell lines obtained by cation-exchange chromatography. The mAb variants produced in the different expression systems are represented as: NS0-R99 (magenta), CHO-R99 (blue), HEK-R99 (black). The profiles show the distribution of acidic and basic species relative to the main peak, with acidic species appearing before the main peak (left side) and basic species appearing after the main peak (right side). Two batches of each variant were analyzed.

**Supplemental Figure 5.**
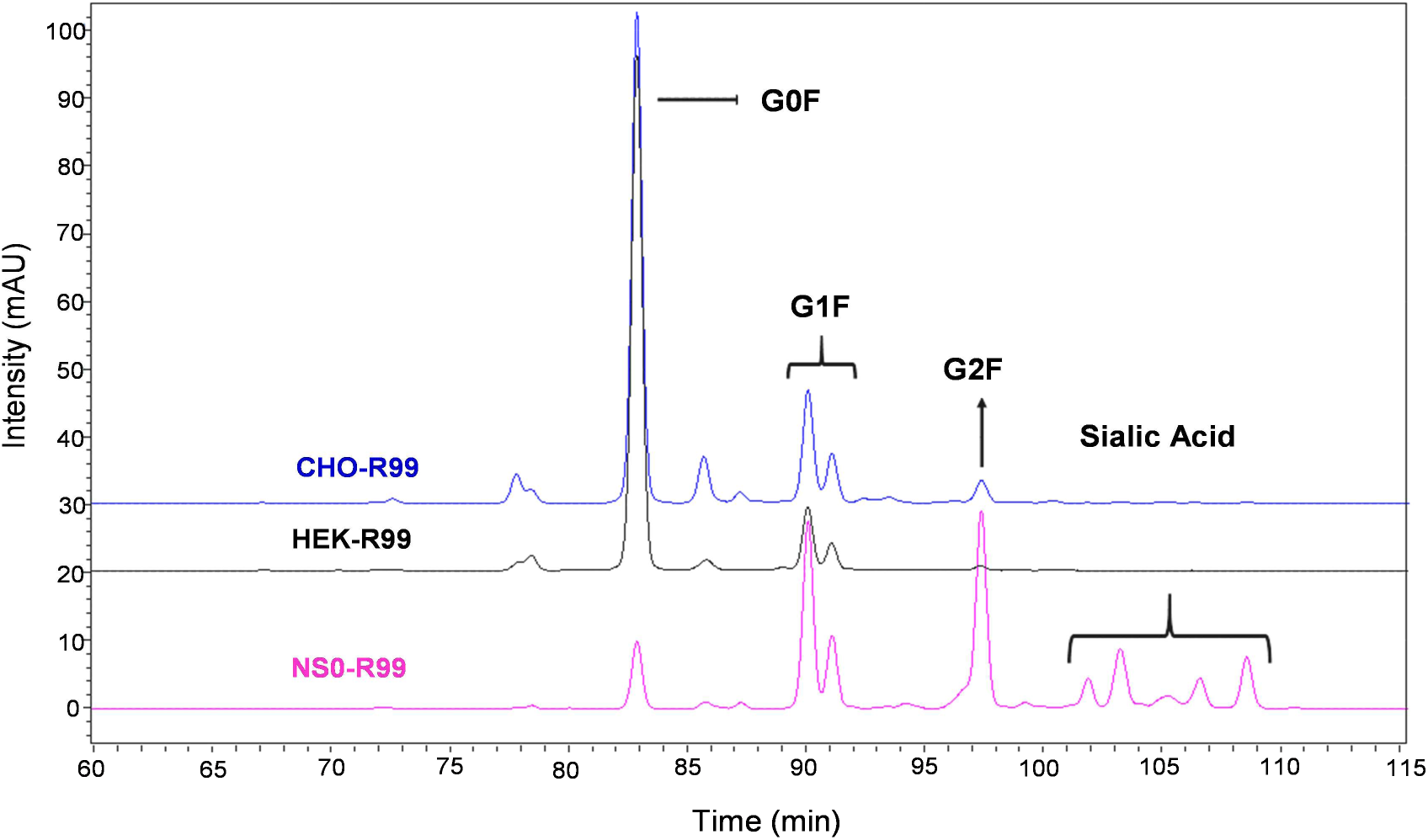
Comparative N-glycosylation profiles of the R99 mAb expressed in NS0, CHO-K1, and HEK-293 host cell lines. The mAb variants produced in the different expression systems are represented as: NS0-R99 (magenta), CHO-R99 (blue), HEK-R99 (black). The profiles display the relative abundance of glycoforms, including G0F (core fucosylated without galactose), G1F (core fucosylated with one galactose), and G2F (core fucosylated with two galactose residues) and sialic acid content, allowing for comparison of the glycosylation patterns across the different expression systems.

